# Seasonal stability and species specificity of environmentally acquired chemical mating signals in orchid bees

**DOI:** 10.1101/2022.09.23.509236

**Authors:** Kathy Darragh, Tess A Linden, Santiago R Ramirez

**Affiliations:** University of California, Davis; Harvard University

**Keywords:** mate choice, signals, pheromone, courtship, reproductive isolation, seasonality

## Abstract

Traits that mediate reproductive isolation between species, such as those involved in mate choice and/or recognition, are predicted to experience stabilizing selection towards the species mean. Male orchid bees collect chemical compounds from many sources, such as plants and fungi, which they use as a perfume signal (pheromone) during courtship display. Environmentally acquired signals are more prone to variation as source availability can vary through space and time. Here, we investigate the seasonality and species-specificity of male perfumes across an entire year in three sympatric species of *Euglossa* orchid bees. Our analysis revealed considerable within-species variation in perfumes. However, species-specificity was maintained consistently throughout the year, suggesting that these perfumes could play an important role in reproductive isolation. Our analysis also identified strong correlations in the abundance of some compounds, possibly due to shared collection sources between species. Our study suggests that orchid bee perfumes are robust in the face of environmental changes in resource availability.

## Introduction

The maintenance of distinct species relies on reproductive isolating barriers that reduce or prevent gene flow between diverging lineages (Coyne and Orr 2004). A key barrier to gene flow in animals is mate choice (Jiggins et al. 2001; West and Kodric-Brown 2015; Martin and Mendelson 2016; Shahandeh et al. 2018). For mate choice to effectively maintain reproductive isolation among closely related lineages, each species must differ in traits associated with mating and/or courtship behavior, and individuals must exhibit a preference for the conspecific phenotype (Mas and Jallon 2005; Ryan and Guerra 2014; Saveer et al. 2014; Shahandeh et al. 2018). Due to their importance in reproductive isolation, traits associated with courtship display and/or mate recognition are expected to experience stabilizing selection, resulting in reduced intraspecific variation and consistent species differences (Gerhardt 1982; Pfennig 1998; Benedict and Bowie 2009; McPeek et al. 2011).

Detection of chemical signals (Robertson 2019) is considered to be the most ancient and widespread sensory system, playing a key role in communication (Ache and Young 2005; Amo and Bonadonna 2018). Of particular relevance to mate choice are sex pheromones: chemical signals that mediate intraspecific communication in the context of mating (Wyatt 2003, 2014). Due to their important role in mating, divergence in signals and preferences between populations can lead to reproductive isolation (Schneider 1992; Johansson and Jones 2007; Smadja and Butlin 2008; Saveer et al. 2014). The role of chemical signaling in speciation has been well-studied in moths, where pheromones experience stabilizing selection towards the species mean (Löfstedt 1993; Smadja and Butlin 2008). However, even with high species-specificity, pheromones exhibit qualitative and quantitative differences within and between populations of the same species which may be due to genetic drift or varying selection pressures either in space or time (Carde and Allison 2016).

The term pheromone refers to the role of the chemical signal but does not address the source of the compound. The mechanisms by which pheromones are acquired, or produced, could impact the amount of intraspecific variation they exhibit depending on the availability and quality of the sources. Some of the most well-studied insect pheromones are biosynthesized *de novo*, for example, many lepidopteran pheromones (Roelofs and Rooney 2003; Liénard et al. 2008; Groot et al. 2016; Darragh et al. 2020, 2021). These genetically controlled pathways could reduce the amount of intraspecific variation due to a lack of reliance on source availability. However, some pheromone compounds are not biosynthesized by the insect itself and instead originate exogenously. For example, arctiid moths, such as *Uthetheisa ornatrix*, sequester alkaloids as larvae which they then process to produce pheromone compounds as adults (Conner et al. 1981).

A unique example of compound acquisition comes from the orchid bees, a group insect pollinators found throughout the lowlands of tropical America, from Mexico to Brazil. Male orchid bees collect compounds from environmental sources, such as flowers and fungi, and store them in specialized hindleg pouches for use as a pheromone (perfume) during mating displays (Vogel 1966; Dressler 1982; Eltz et al. 1999, 2005b). In addition to the perfume compounds that orchid bees collect, male bees accidentally incorporate many additional by-product compounds that co-occur with the compounds they actively search for (Eltz et al. 2005a). These additional compounds may vary between individuals as male bees collect perfume compounds from multiple sources (Ramírez et al. 2002; Pemberton and Wheeler 2006). Due to the reliance of orchid bees on environmental sources these signals could be prone to exhibiting a substantial amount of variation across both space and time.

The stability and species-specificity of orchid bee perfumes has mainly been investigated with respect to geography. Orchid bees can be found in areas with differing plant communities. For example, an introduced population of *Euglossa dilemma* in Florida, a region lacking perfume orchids, has a high level of perfume similarity compared to bees from the native range in Mexico and Central America (Pemberton and Wheeler 2006; Ramírez et al. 2010a). *Euglossa dilemma* and *Euglossa viridissima*, a recently diverged pair of orchid bees, exhibit consistent species-specific differences across their ranges (Brand et al. 2020). Moreover, these perfume difference coincide with rapid evolution of odorant receptor genes that mediate both perfume acquisition by males and perfume preference by females, resulting in reproductive isolation (Brand et al. 2020). Comparisons across more distantly related lineages have also found evidence for species-specificity of perfumes, with much greater variation between species than within species (Zimmermann et al. 2006; Weber et al. 2016).

In addition to variation in space, orchid bee perfumes could vary in time. The availability of chemical compounds may change throughout the year as source abundance fluctuates due to phenological cycles. Although orchid flowers provide only a fraction of the compounds collected by orchid bees in their perfumes (Whitten et al. 1993; Ramírez et al. 2011), many orchid species exhibit a pronounced flowering peak in the dry season in Panama, with few species exhibiting year-round blooming patterns (Ackerman 1983). A peak in orchid diversity within bee-orchid interaction networks also occurs during the dry season in Costa Rica (Ramírez 2019). However, relatively little is known about orchid bee perfume dynamics during these seasonal changes. Studies comparing one timepoint per season, find mixed evidence of seasonal effects. *Euglossa dilemma* has a more complex perfume in the rainy season, but only marginal effects on complexity are seen in *Euglossa viridissima* (Eltz et al. 2015). The same dataset did not find seasonality of individual compounds (Pokorny et al. 2013). However, these studies of two timepoints do not represent a true time series.

Here, we conducted a year-long analysis of perfume variation in three co-occurring species of orchid bees. We analyze perfume composition of 572 individual male bees from two closely related species, *Euglossa imperialis* and *Euglossa flammea*, and a more distantly related euglossine bee, *Euglossa tridentata* (Ramírez et al. 2010b). Samples were collected at monthly intervals over a year, resulting in a time series dataset which we use to study the seasonality of orchid bee perfumes. We describe how species differ in their perfumes, which compounds contribute to these differences, and how consistent these differences are through time. We also carry out intraspecific analyses to investigate the seasonality of the perfume of each species and whether compound collection exhibits seasonal trends.

## Methods

### Sample collection

Samples were collected in La Gamba field station, Puntarena, Costa Rica (8°42′03′’ N, 83°12′06′’ W) from 28^th^ August 2015 (referred to as September 2015 samples), until 30^th^ August 2016 (referred to as September 2016 samples) between 8am and 12pm. Samples were collected at approximately one-month intervals (exact dates found in sample information at https://osf.io/rwxv6/?view_only=1851fc1b29de4b8eaaf1f0657d9ee876). For most analyses these timepoints were considered separately, however, for seasonality analyses where 12 timepoints (one per month) are required, we combined samples from September 2015 and September 2016. Bees were collected by netting at chemical baits on filter paper using cineole, eugenol, and methyl salicylate. Precipitation data is available from La Gamba field station (https://www.lagamba.at/en/research/scientific-data-of-the-golfo-dulce-region/).

### Chemical analysis

Hindlegs were placed in 500μl hexane and stored at *20°C. For analysis, 50μl was transferred to a vial containing 15μl of a 16.5ng/μl solution of 2-undecanone in hexane as an internal standard. Samples were analyzed using Agilent model 5977A mass-selective detector connected to Agilent GC model 7890B, with a HP-5 Ultra Inert column (Agilent, 30 m × 0.25 mm, 0.25 µm). 1μl of each sample was injected using Agilent ALS 7694 autosampler in split mode with a 5:1 ratio with helium as the carrier gas (250°C injector temperature, split flow of 3.5 ml/min). The temperature program started at 55°C for 3 minutes, and then rose at 10°C/min to 300°C. The temperature was held at 300°C for 1 minute and 315°C for 5 minutes.

Compounds were identified by comparing mass spectra and gas chromatographic retention index with previous analyses. Compounds not thought to be perfume compounds, such as hydrocarbons or compounds also found in head extracts, were removed. Many are likely to be derived from labial gland compounds which the male bees release to dissolve volatiles before transferring this mixture to the hindlegs and recycling the labial compounds (Eltz et al. 2007). We included volatile/semi-volatile compounds eluting before a retention index of 2400. We removed compounds found in less than five percent of individuals from the overall dataset and repeated this when analyzing data from each species.

### Statistical analyses

#### Do species differ in their perfumes?

To measure perfume divergence, we carried out nonmetric multidimensional scaling (NMDS) (Bray-Curtis similarity matrix, lowest k value with stress<0.2 was k=4) using the “metaMDS” function in *vegan* with absolute peak areas (Oksanen et al. 2020). For visualization we used the *ade4* package (Dray and Dufour 2007; Thioulouse et al. 2018).

We used multivariate analyses to investigate perfume variation. We carried out a PERMANOVA (permutational multivariate analysis of variance) using the “adonis2” function in *vegan* (Bray-Curtis distance matrix, 1000 permutations). We tested each term sequentially, starting with species, as this was the main clustering factor identified through visualization, followed by month, and an interaction term. To evaluate model fit, we used Akaike’s information criterion (AIC)(Table S1). To identify which groups were significantly different from each other we carried out Bonferroni-corrected *post hoc* pairwise testing using the “pairwise.perm.MANOVA” function in the *RVAideMemoire* package (Hervé 2021).

Distance-based analyses can lead to false-positives by confounding differences in dispersion and location (Warton et al. 2012). We tested for differences in variance using the “betadisper” and “permutest” functions in *vegan*. To confirm the results of the PERMANOVA analysis, we used multivariate generalized linear models using the function “ManyGLM” from the *mvabund* package (Wang et al. 2012). We rounded our data to integers and modelled using a negative-binomial distribution. The “ManyGLM” function fits models to each compound in the dataset and then sums the test statistics producing a multivariate test statistic known as Sum-of-LR, which can be tested for significance using resampling. We included species, month, and an interaction term. We used backward elimination and compared model fit with a likelihood ratio test (Table S2). The output includes the contribution of each compound to the Sum-of-LR, allowing us to determine which compounds drive group differences. P-values were adjusted for multiple testing.

#### Which compounds contribute to these species differences?

In addition to identifying the compounds driving group differences using ManyGLM, we also carried out an indicator analysis using the *indicspecies* package to determine which compounds contribute to species differences (Cáceres and Legendre 2009). The groups of interest are the species, and the goal is to identify compounds which indicate group membership. The best indicator would be a compound which is found in a single species (specificity) and in all members of that species (coverage), resulting in a perfect indicator value of one. Compound specificity is calculated using amounts, while coverage only includes presence/absence data. We used the function “multipatt” to investigate which single compounds are the best predictors of membership to each species (De Cáceres et al. 2012).

#### Do species share perfume motifs?

It has been suggested that closely-correlated compounds are likely derived from the same perfume sources (Zimmermann et al. 2009). To determine if the species in our analysis shared groups of correlated compounds, we created correlation matrices using the “cor” function in the *corrplot* package (Wei and Simko 2021). We tested for significant correlations using the “cor.mtest” function. We plotted the significant strong correlations (with a cut-off of p=0.01 and R<0.8) using hierarchical clustering in the “corrplot” function and compared clusters between species.

#### Are species differences consistent through time?

To visualize differentiation between species throughout the year we calculated Bray-Curtis differences in a pairwise fashion each month and plotted the resulting differences to show how average species differences change over time.

We conducted statistical analyses to determine how species differences change over time. The dynamics of a particular species over time can be considered as a trajectory through space using community trajectory analysis (De Cáceres et al. 2019; Sturbois et al. 2021). We reduced each time point to the average compound amount for all compounds for each species so that each month only has one multivariate datapoint per species. We used the function “trajectoryPCoA” from the package *ecotraj* to display the trajectories for each species. To investigate the geometric properties of each trajectory, we used the functions “trajectoryLengths” and “trajectoryDirectionality” to determine trajectory length and directionality. To compare trajectories between species we used the functions “trajectoryDistances” to calculate the average distance between each species, and “trajectoryConvergence” to test for convergence between species over time.

In this analysis we assume that species would either converge or diverge over time, however, species differences could vary seasonally. To test this, we calculated the centroid of all individuals of each species per month in the NMDS ordination space. For each month, we then calculated the Euclidean distance between cluster centroids (using all 4 NMDS axis) resulting in one distance value for each species-comparison per month (McLean et al. 2019). For each species-pair comparison we then used the “cosinor” function in the *season* package (Barnett and Dobson 2010; Barnett et al. 2012, 2021). This function fits a cosinor model as part of a generalized linear regression, assuming a sinusoidal pattern of seasonality. We log-transformed our data and used the gaussian distribution, found to be appropriate based on residual plots. We assumed that one cycle occurs per year, with one peak and one trough, explained by the phase of the model. Each cosinor model has two terms, sine and cosine, which define the sinusoid and have associated *p*-values. The threshold for significance is reduced to 0.025 to account for multiple testing. We also corrected for multiple testing due top number of compounds using the “p.adjust” function in R with the false discovery rate option.

#### Does compound collection exhibit seasonality?

In addition to testing whether overall species differences exhibit seasonality, we wanted to investigate whether compound collection within species exhibits seasonality. Including month in the PERMANOVA and ManyGLM models tests whether compounds change over time, however, this ignores the order of the months, instead of including the likely correlation between consecutive months. To account for this correlation we used the “cosinor” function in the *season* package (Barnett and Dobson 2010; Barnett et al. 2012, 2021), assuming one cycle per year. We did this both for the amount of each individual compound collected by a species throughout the year, and for NMDS dimensions for each species. The NMDS analyses were run for each species (k=2 *E. flammea* and E. *imperialis*, k=3 *E. tridentata*, lowest k value with stress<0.2 chosen). We log-transformed our data (+2 to allow us to log the negative values from NMDS dimension scores and +1 to allow us to log the zero values for the individual compounds) and used the gaussian distribution, found to be appropriate based on residual plots. As above, the significance threshold is reduced to 0.025 to account for multiple testing. We corrected for multiple testing across multiple compounds per species using the “p.adjust” function R with the false discovery rate option.

#### Plotting and data manipulation

Plots were made using *ggpubr* (Kassambara 2019), *cowplot* (Wilke 2020), and *ggplot2* (Wickham 2009). Additional packages used for data transformation were *MASS* (Venables et al. 2002), *dplyr* (Wickham et al. 2021), *tibble* (Müller and Wickham 2022), and *usedist* (Bittinger 2020). Analyses were carried out in R version 4.1.2 (R Core Team 2021).

## Results

### Do species differ in their perfumes?

We sampled 572 male orchid bees of three species across one year (12-16 individuals per species per month) and identified 222 compounds. All species differed both in the total number of compounds, and the total amount of compound present in their perfume (Figure S1). Overall, *E. tridentata* had both the highest number of compounds and the largest quantities of the combined compounds. While there was some overlap in the compounds found in each species, the most abundant compounds differed considerably (Table 1).

**Table 1.** Five most abundant compounds in each orchid bee species (percentage of total perfume).

| *E. flammea* | *E. imperialis* | *E. tridentata* |
| --- | --- | --- |
| (Z)-carvone oxide 52% | cineole 30% | (E)-β-ocimene 12.7% |
| carvone 5.4% | germacrene d 16.6% | 2,3-epoxygeranylacetate 7.7% |
| 2-methylformalinide 4.3% | hexahydrofarnesylacetone 13.3% | eugenol 7.3% |
| (E)-limonene oxide 4.3% | nerolidol 4.8% | 4-methoxycinnamylalcohol 6.3% |
| cineole 4.1% | α-phellandrene 2.7% | geranylgeraniol 6.1% |

Both *E. flammea* and *E. imperialis* have simpler perfumes, dominated by one or a few compounds, in contrast to the more diverse perfume of *E. tridentata* (Figure S2). The perfume of *E. flammea* is the simplest, with (*Z*)-Carvone oxide averaging 52% of the perfume (Table 1). The perfume of *E. tridentata* is more complex, and includes many low-abundance compounds; the most abundant compound is only 12.7% of the total perfume (Table 1). The most abundant compounds in *E. flammea* are also found in 99% of individuals, showing that these are the primary focus of male collection (Table 2). In contrast, the compound with highest frequency in *E. tridentata*, (*Z*)-linalool oxide, found in 91% of individuals, is not included among the five most abundant compounds (Table 2). In general, the frequency of compounds shows a different pattern to compound abundance since many compounds that are found at high frequency are not present in high abundance (Figs. S2, S3).

**Table 2.** Five compounds most frequently identified in each orchid bee species (percentage of bees containing compound)

| *E. flammea* | *E. imperialis* | *E. tridentata* |
| --- | --- | --- |
| (Z)-carvone oxide 99% | hexahydrofarnesylacetone 97% | (Z)-linalool oxide 91% |
| 2-methylformalinide 99% | cineole 95% | 2,3-epoxygeranylacetate 90% |
| carvone 97% | germacrene d 95% | (E)-β-ocimene 83% |
| cineole 96% | nerolidol 94% | unknown RI=1318.1 79% |
| nerolidol 95% | δ-cadinene 92% | 1,4-dimethoxybenzene 79% |

To determine the species-specificity of perfumes, we investigated how variation is partitioned between and within species. Individuals mostly cluster by species (Fig. 1). Species have significantly different perfumes, with species accounting for 37% of variation in perfume (PERMANOVA *F*_2,571_ = 174.28, *p* < .001). All pairwise comparisons of species are significantly different (Bonferroni-corrected pairwise PERMANOVA, p=0.003). A further 3% of the variation is explained by collection month (PERMANOVA, F_12,571_ = 2.29, *p* < 0.001). Since species also differed in their dispersion (permutation test of homogeneity of dispersion, F_2,569_=19.86, p=0.001; Table S3) we confirmed these results with multivariate generalized linear models using the *mvabund* package (Wang et al. 2012; Warton et al. 2012). We found the best model included species and month, with more variation explained by species, as detected by PERMANOVA (Table S4).

**FIGURE 1.**
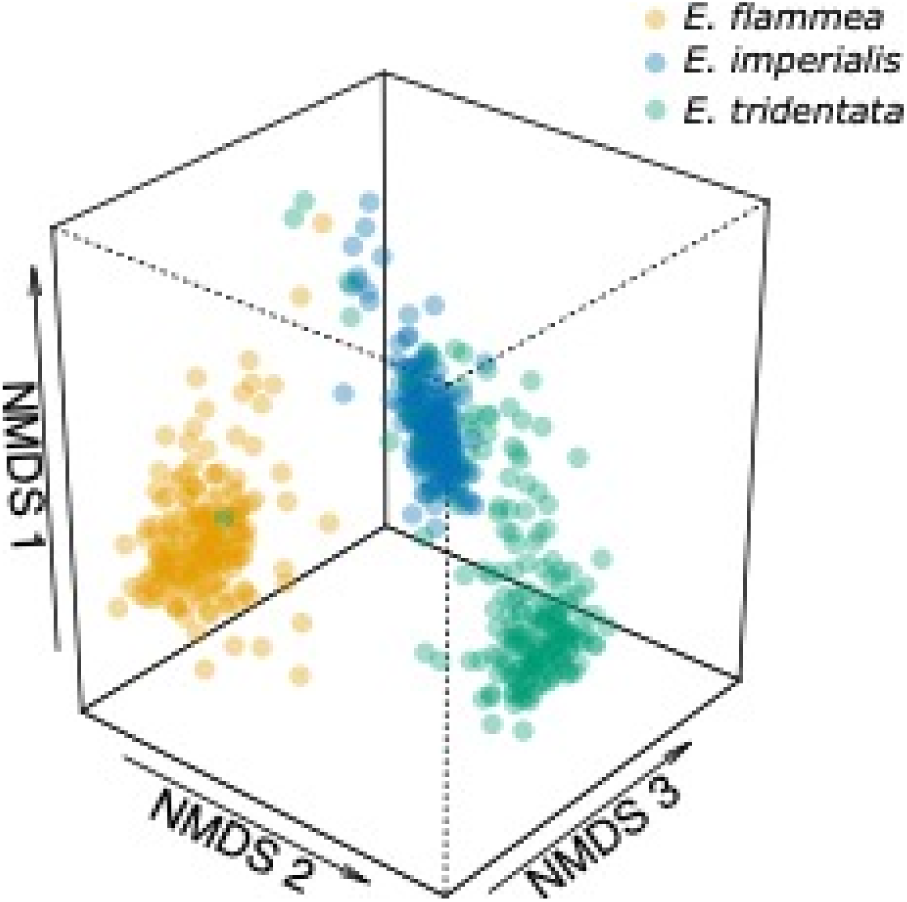
NMDS (nonmetric multidimensional scaling) plot illustrating in three dimensions the variation in the perfumes of males of three *Euglossa* species: *E. flammea, E. imperialis*, and *E. tridentata*. Stress=0.09.

### Which compounds contribute to species differences?

To determine which compounds best predict membership to a particular species we carried out an indicator analysis. The best predictors of species identity are those which are found in every individual of a species and in no individuals of any other species. Therefore, it is not always the case that the most abundant compound in a species in the best indicator as it may also be found in other species. For example, cineole is the most abundant compound in *E. imperialis* but is also found in *E. flammea*, making it a poor predictor of species identity. We found that 2/3 of indicator compounds in *E. flammea* and *E. tridentata*, and 1/3 in *E. imperialis* were also in the top five most abundant compounds for those species (Table 1, Table 3). Some of these compounds were also found to be major contributors to deviance due to the species term in the ManyGLM model (Table S4).

**Table 3.**
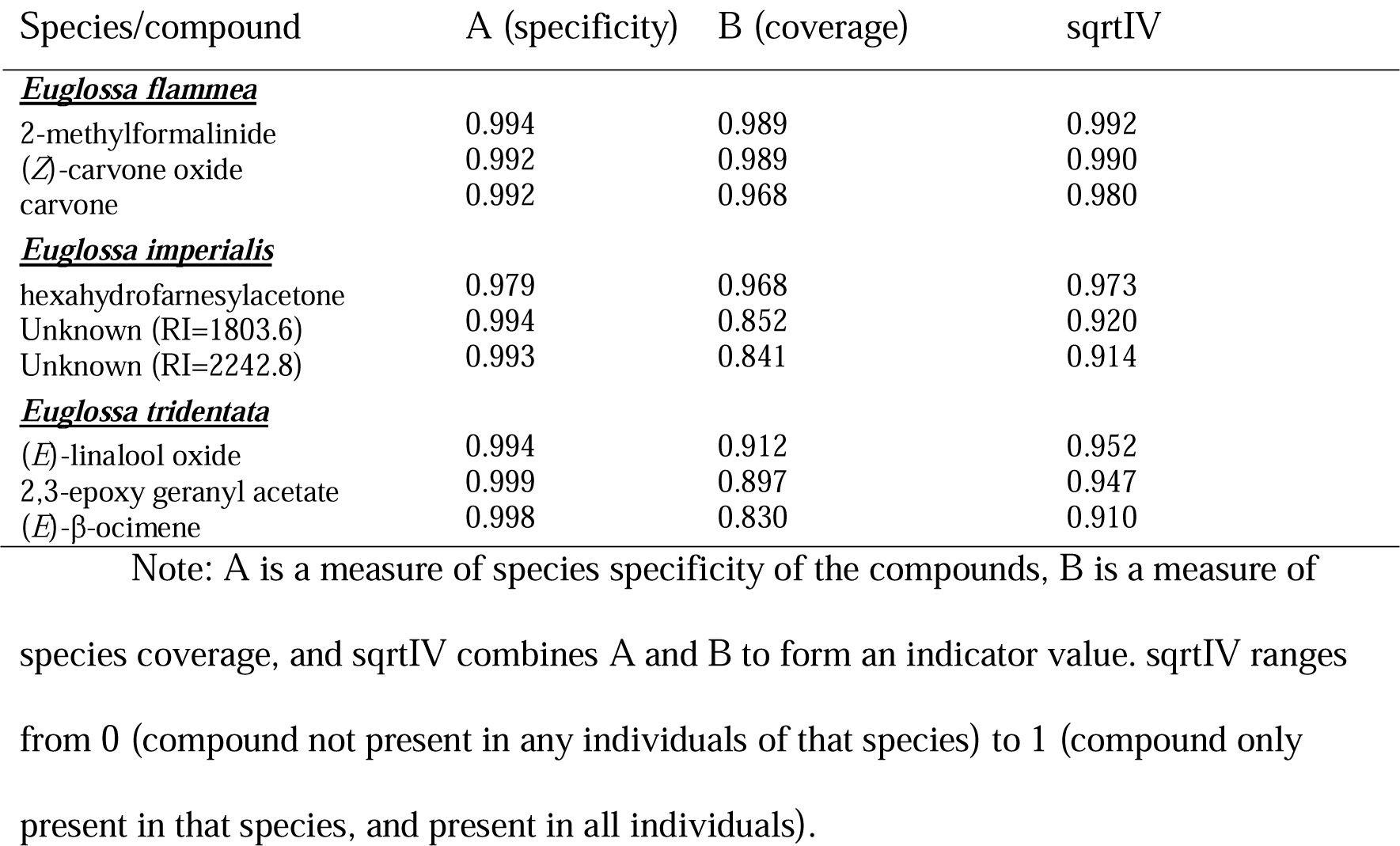
Compounds which are the best indicators of species identity.

### Do species share perfume motifs?

It has been suggested that closely correlated compounds (known as motifs) are likely to be derived from the same perfume source (Zimmermann et al. 2009). This implies that motifs shared among individuals of the same species (or different species) correspond to compounds obtained from the same perfume sources. To test this, we calculated inter-compound correlation within each species. Overall, as expected due to the fact that orchid bees use a diverse range of sources for collection, we found that most compounds vary independently, with a low level of correlation between compounds (*E. imperialis, R*=0.09; *E. tridentata, R*=0.1; *E. flammea, R*=0.14). The biggest motif found in *E. imperialis* is formed of eight sesquiterpenes and similar to a six-compound motif found in *E. flammea* (Figs. S4, S5). Another motif, this time of acetates, is also shared between *E. imperialis* and *E. flammea*. In addition, *E. flammea* has a species-specific motif consisting mostly of carvone and limonene compounds (Fig. S2). The main motifs identified in *E. tridentata* are smaller and generally not shared with the other species (Fig S4). Some motifs made up of only two compounds were shared between all three species such as α-terpinene and γ-terpinene (Figs. S4, S5, S6).

### Are species differences consistent through time?

Visualization of species differences through time revealed that interspecific differences are maintained throughout the year for all three species pairs (Figure 2). We used community trajectory analysis to track the trajectory of each species through time in our study period. We found that while *E. flammea* and *E. tridentata* have similar trajectory lengths, meaning change in perfume composition between months, *E. imperialis* has a trajectory length of less than one third of the other two species (Figure S7). *Euglossa flammea* changes most over PCoA1 which accounts for a higher percentage of variation suggesting that this species exhibits the biggest changes. All three species exhibit low levels of directionality, suggesting little overall change in perfume composition through time (Fig. S7). Similar to our NMDS visualization, we found that *E. flammea* and *E. tridentata* were the most dissimilar (average distance between trajectories: *E. flammea* – *E. tridentata*, 110,750; *E. flammea* – *E. imperialis*, 94,116; *E. imperialis* – *E. tridentata*, 86,438). Finally, we found no evidence for convergence or divergence in chemical similarity between species (Mann-Kendall trend test, p=NS). We followed up this linear analysis with a seasonality analysis where species differences through the year are modelled as a sinusoidal curve. We found no evidence for seasonal changes in species differences throughout the year (Table S5).

**FIGURE 2.**
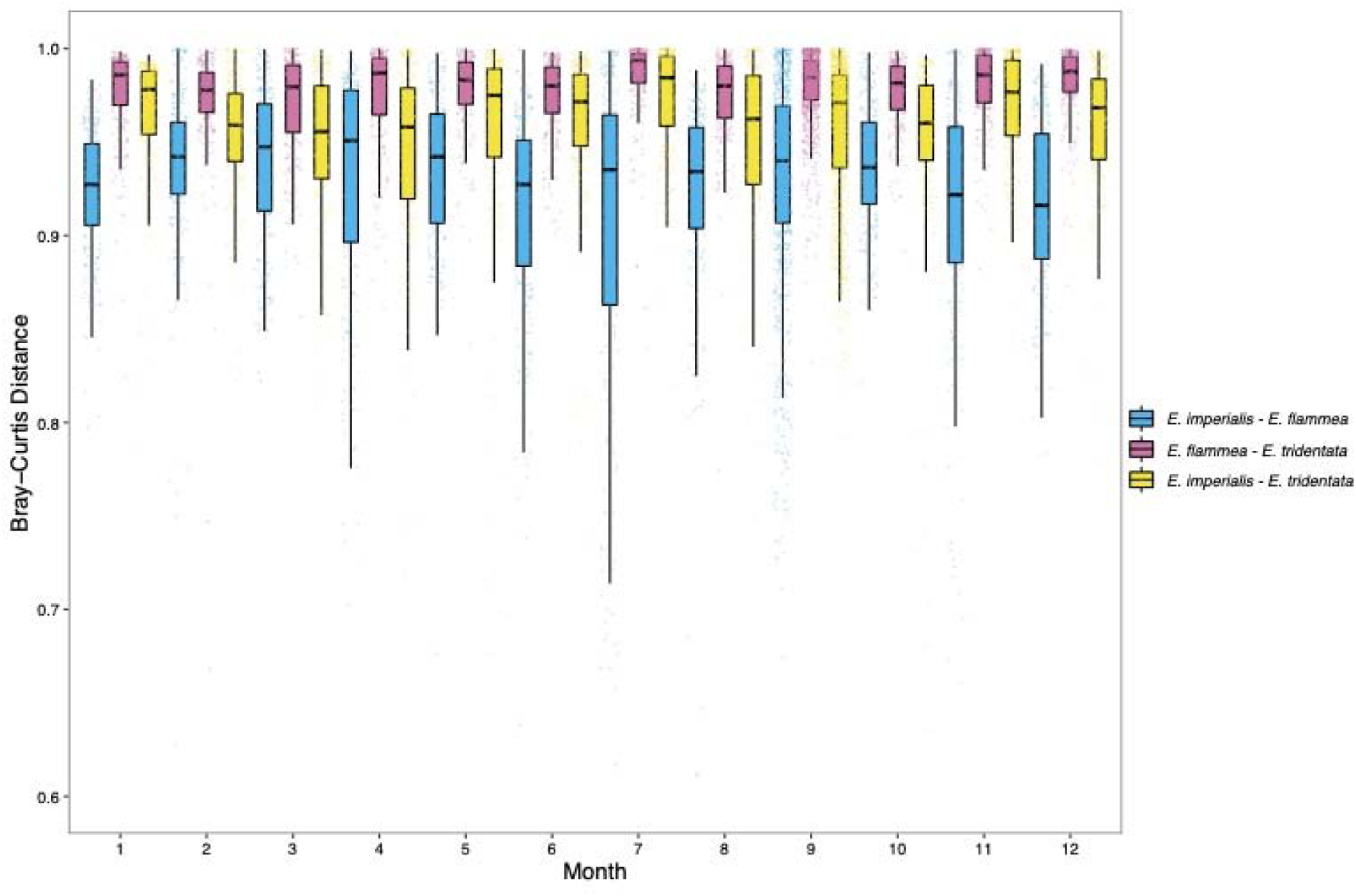
Pairwise Bray-Curtis distances between *E. imperialis, E. flammea* and *E. tridentata* for each month of the year. More data points are included in Month 9 as this month was sampled in two different years at the start and end of sampling. For the x-axis, 1 is January and 12 is December. 23 outlier comparisons were removed with low Bray-Curtis distances.

### `Does compound collection exhibit seasonality?

To test whether compound collection exhibits seasonality, we used cosinor model analyses. Firstly, we took a multivariate approach by looking for seasonal patterns in the NMDS ordinations of each species. We found seasonal effects for the first NMDS dimension of both *E. flammea* and *E. tridentata*, as well as the second NMDS dimension of *E. tridentata*, while no dimension in *E. imperialis* exhibited seasonal variation (Table S6).

We then tested individual compounds for evidence of seasonality. We found that 39% of *E. flammea* compounds (41/105), 35% of *E. imperialis* compounds (48/139), and 22% of *E. tridentata* compounds (40/184) exhibit a pattern of seasonality. The seasonal compounds found in each species are not mutually exclusive, with eight shared between all three species (RI=1203.5, ethyl,4-ethoxybenzoate, cineole, geranyl linalool, α-terpineol, α-phellandrene, RI=1081.5 (acetate), and phenyl acetaldehyde). A similar peak phase across species was found for most compounds, suggesting that seasonality could be due to environmental abundance of the compounds. Despite compound seasonality, species differences are maintained throughout the season, for example, cineole is always found in higher absolute and relative abundance in *E. imperialis* even during seasonal fluctuations (Fig. 3). Of the ten compounds which contributed most to the deviance explained by “month” in the Many GLM model, seven were also identified as seasonal compounds, with four identified as seasonal in all three species. We found that fewer compounds exhibit a pattern of seasonality when analyzing relative abundance: 21% of *E. flammea* compounds (23/105), 12% of *E. imperialis* compounds (16/139), and 13% of *E. tridentata* compounds (23/184) exhibit seasonality. Full results table of all compounds for each species is found on the OSF (https://osf.io/rwxv6/?view_only=1851fc1b29de4b8eaaf1f0657d9ee876).

**Figure 3.**
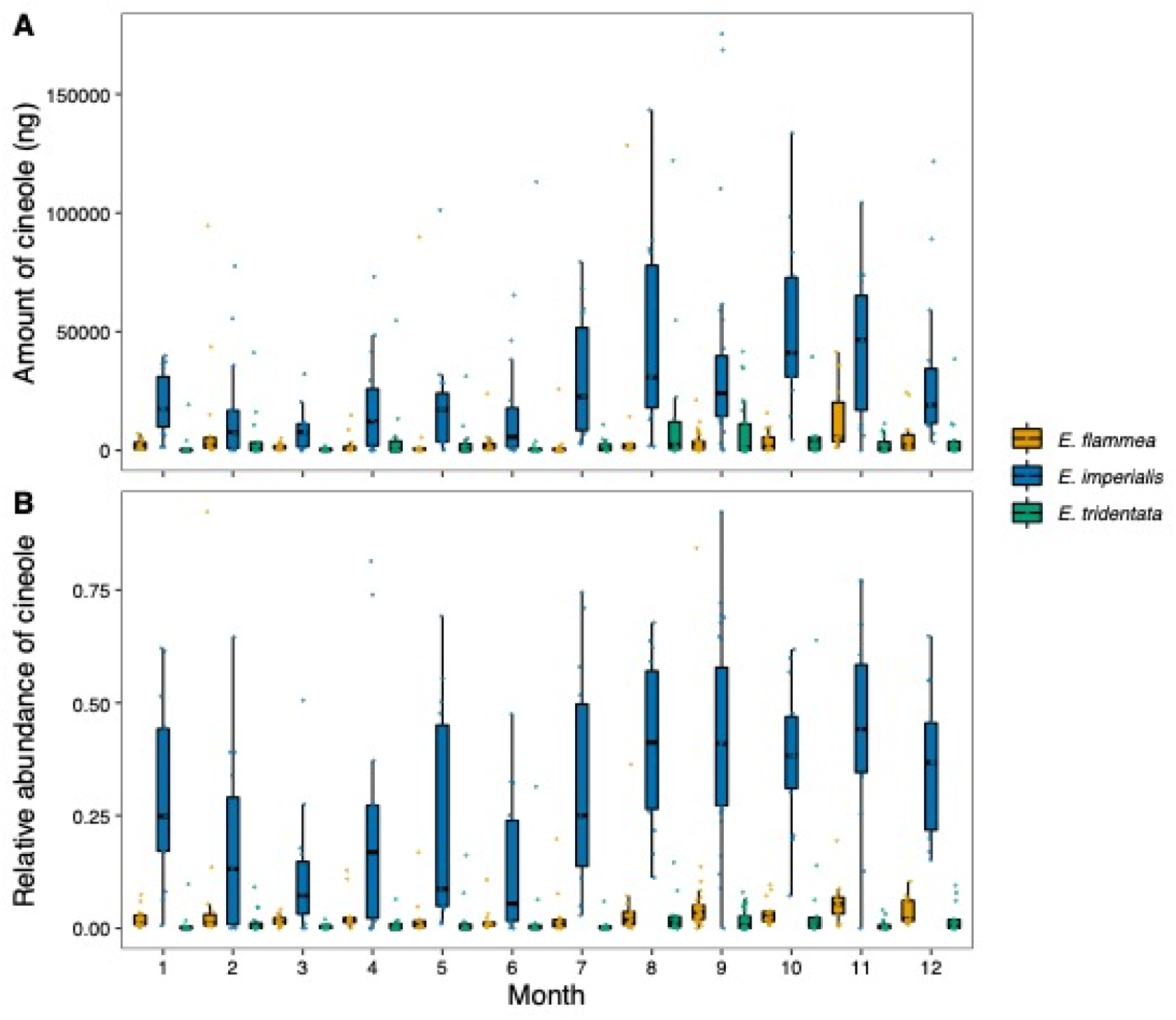
(a) Absolute amount of cineole in male *E. imperialis, E. flammea* and *E. tridentata* for each month of the year. (b) Relative abundance of cineole in male *E. imperialis, E. flammea* and *E. tridentata* for each month of the year. More data points are included in Month 9 as this month was sampled in two different years at the start and end of sampling. For the x-axis, 1 is January and 12 is December.

These trends are not just due to overall increases or decreases in collection throughout the year. We found no evidence for seasonality in number or total amount of compounds collected, except for the amount of compound collected by *E. tridentata* which peaked in June (Table S7). In addition, we did not find a correlation between compound abundance or frequency and seasonality. Seasonal compounds do not differ in their mean abundance relative to non-seasonal compounds (ANOVA: *E. flammea*, F_1,103_=2.34, *p*=NS; *E. imperialis*, F_1,137_=0.22, *p*=NS; *E. tridentata*, F_1,182_=0, *p*=NS). Seasonal compounds also do not differ in their frequency relative to non-seasonal compounds (ANOVA: *E. flammea*, F_1,103_=0.076, *p*=NS; *E. imperialis*, F_1,137_=0.59, *p*=NS; *E. tridentata*, F_1,182_=0.001, *p*=NS).

We found that all three species exhibited similar seasonality in their compound collection. We looked at the peak month for all compounds identified as seasonal in each species and found no differences in mean peak collection month (Fig. 4). The mean for *E. flammea* was found in late May (phase=5.8), while for *E. imperialis* and *E. tridentata*, the mean was mid-June (phase=6.5 and phase=6.3, respectively). While there was no difference between the mean peak month for compound seasonality in each species, violin plots show that the distribution differs. *E. imperialis* and *E. tridentata* have most peaks in the early-mid rainy season (Fig. 4), while *E. flammea* has a more even spread throughout the year. We found no difference in peak phases between absolute and relative analyses (Figure S8).

**FIGURE 4.**
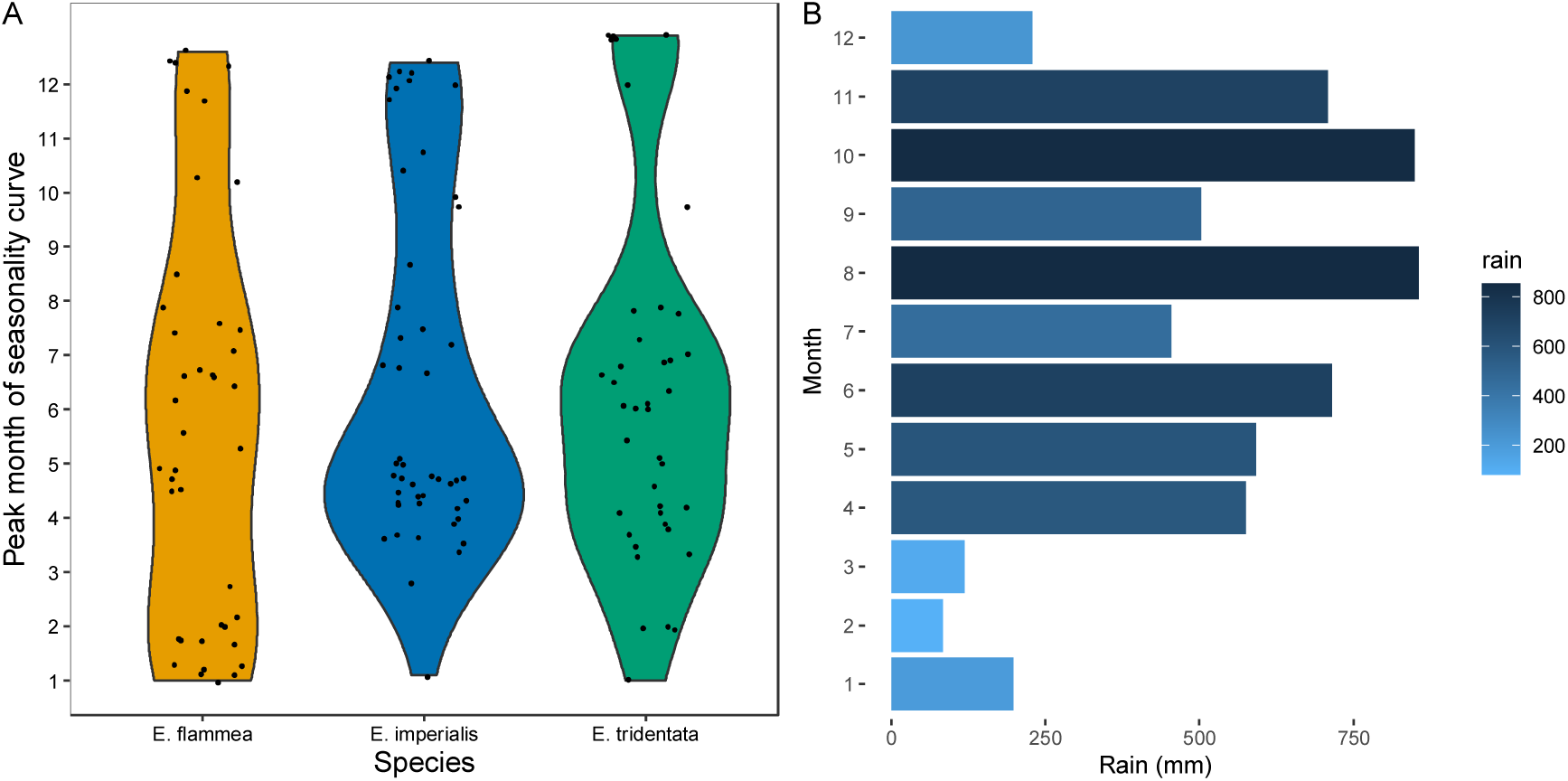
(a) Violin plot illustrating the variation in the peak month of the seasonality curve for compounds of each species. Only compounds which were determined to exhibit seasonality were included. Species did not differ in their peak month of seasonality (Kruskal Wallis, df=2, H test statistic=0.83, p=NS). (b) Rain data from La Gamba field station in the years 2015-2016. Data from September 2015 and 2016 was combined and the average taken. For the y-axes, 1 is January and 12 is December.

## Discussion

The unique nature of orchid bee perfume collection makes it an excellent example to study the dynamics of chemical communication. Male orchid bees collect chemical compounds from a range of exogenous sources which they use as perfumes during courtship. Here, we investigate the chemical ecology of three sympatric species of orchid bee, testing how species differ in their perfumes and whether these environmentally derived mating signals exhibit seasonality. We found that, as previously described, orchid bees exhibit high levels of species specificity in their perfumes. We show that species differences are maintained over time with remarkable consistency throughout the year. Differentiation between species is maintained despite intraspecific variation, seasonality in compound collection, and potentially shared collection sources between species. Our results suggest an astounding robustness of orchid bee perfume chemical signals in the face of changing environmental conditions and available resources even though male bees rely exclusively on exogenous sources for perfume formation.

Orchid bee perfumes exhibit remarkable species specificity and remain stable across a large geographic range (Zimmermann et al. 2006; Ramírez et al. 2010a; Weber et al. 2016). We find that this species specificity is also maintained through time, with species maintaining consistent differences throughout changing seasons. In comparison with species which biosynthesize their pheromones, such as *Heliconius* butterflies, we find more intraspecific variation in orchid bees (variation explained by species: *Heliconius*, 58%; orchid bees, 37%) (Darragh et al. 2017, 2019, 2020). Nonetheless, species identity is the best predictor of perfume divergence among orchid bees, with greater interspecific variation than intraspecific variation. The pattern of species-specificity and consistency detected suggests that orchid bee perfumes are under strong stabilizing selection, as predicted for signals important for reproductive isolation (Löfstedt 1993).

We find that most variation in orchid bee perfumes reflects species identity and not local resource availability. Orchids, and the majority of plants, have been described to flower during the dry season in the tropical forests of Central America (Fournier and Salas 1966; Janzen 1967; Croat 1969; Frankie et al. 1974; Ackerman 1983; Ramírez 2019). However, male bees do not only rely on floral sources alone for perfume collection and have been described to collect compounds from many types of sources, including rotten or fungus-infected logs, exposed plant roots, leaves, and even walls sprayed with insecticide (Roberts et al. 1982; Whitten et al. 1993; Ramírez et al. 2002; Cappellari and Harter-Marques 2010). Male orchid bees search for chemical compounds rather than compound sources, meaning they can switch easily between sources throughout the seasons to fulfill their species-specific preferences. Furthermore, male bees have been proposed to exhibit learned avoidance through negative feedback whereby collection of a particular chemical compound reduces its attractiveness, preventing overcollection (Eltz et al. 2005a; Pokorny et al. 2013). A diversity of perfume sources, alongside a learning mechanism, could buffer orchid bee perfumes from changing due to seasonal conditions.

Many chemical compounds are collected by an individual male orchid bee, making it difficult to determine which compounds are used as perfume and collected purposefully and which are “noise” compounds (Ramírez et al. 2010a). In general, only one or a few compounds are collected in high abundance by each species (Eltz et al. 1999; Zimmermann et al. 2009). In this study, the perfumes of *E. flammea* and *E. imperialis* are dominated by a small number of compounds, while *E. tridentata* has a less clear dominance pattern. We cannot assume that compound abundance is the same as biological importance. We found that many more compounds were found at a high frequency than at high abundance. These compounds may be target compounds for the bees explaining their high frequency, or alternatively could be compounds produced by the perfume sources of the bees, collected as by-products. Ultimately behavioral experiments are needed to determine functionality.

Individual male orchid bees form complex perfumes by collecting compounds from a variety of different sources. It has been suggested that this results in subsets of compounds (“motifs”) which are derived from the same source and are intercorrelated (Zimmermann et al. 2009). We find some overlap with previously identified motifs. We find a motif of short chain acetates previously identified in *E. imperialis* (Zimmermann et al. 2009). However, we do not detect a hexahydrofarnesyl acetone motif, perhaps expected as a widespread compound like this is likely collected from different sources throughout the year, eroding any correlations found in a certain season (Zimmermann et al. 2009). Interestingly, we find shared sesquiterpene and acetate motifs between the closely related *E. imperialis* and *E. flammea*. This could indicate the use of shared compound sources, implying that closely related species can maintain species-specific perfume blends despite sharing compound sources. However, it could also be related to compound synthesis, with compounds originating from the same biosynthetic pathway more likely to be correlated, irrespective of the compound source. Many correlations are between biosynthetically similar compounds such as aromatics, acetates, sesquiterpenes, or even isomers. This might suggest that species have shared motifs due to biosynthetic constraints rather than shared compound sources.

Our study reveals temporal patterns in orchid bee perfumes which are consistent with a role in reproductive isolation between species. Our data revealed strong differences between species that remain consistent throughout the seasons, as well as the presence of species-specific compounds. These findings suggest that orchid bee perfumes are experiencing stabilizing selection towards a species mean. We also find evidence for intraspecific variation and seasonality in the collection of some compounds, perhaps to some extent due to environmental noise through the seasons. Finally, we show that the same highly intercorrelated compounds can be found in multiple species, perhaps indicating the use of shared compound sources. One caveat is that it is unclear which compounds are biologically important for mate choice, with future behavioral studies needed to disentangle this.

## References

Ache, B., and J. Young. 2005. Olfaction: Diverse Species, Conserved Principles. Neuron 48:417–30.

Ackerman, J. D. 1983. Specificity and mutual dependency of the orchid-euglossine bee interaction. Biol. J. Linn. Soc. 20:301–314.

Amo, L., and F. Bonadonna. 2018. Editorial: The Importance of Olfaction in Intra- and Interspecific Communication. Front. Ecol. Evol. 6.

Barnett, A. G., P. Baker, and A. J. Dobson. 2012. Analysing Seasonal Data. R J. 4:5–10.

Barnett, A. G., P. J. Baker, and A. J. Dobson. 2021. season: Analysing Seasonal Data R Functions. R package version 0.3.13.

Barnett, A. G., and A. J. Dobson. 2010. Analysing Seasonal Health Data. Springer Berlin Heidelberg.

Benedict, L., and R. C. K. Bowie. 2009. Macrogeographical variation in the song of a widely distributed African warbler. Biol. Lett. 5:484–487.

Bittinger, K. 2020. usedist: Distance Matrix Utilities. R package version 0.4.0.

Brand, P., I. A. Hinojosa-Díaz, R. Ayala, M. Daigle, C. L. Y. Obiols, T. Eltz, and S. R. Ramírez. 2020. The evolution of sexual signaling is linked to odorant receptor tuning in perfume-collecting orchid bees. Nat. Commun. 11:1–11.

Cáceres, M. D., and P. Legendre. 2009. Associations between species and groups of sites: indices and statistical inference. Ecology 90:3566–3574.

Cappellari, S. C., and B. Harter-Marques. 2010. First Report of Scent Collection by Male Orchid Bees (Hymenoptera: Apidae: Euglossini) from Terrestrial Mushrooms. J. Kans. Entomol. Soc. 83:264–266.

Carde, R. T., and J. D. Allison. 2016. Variation in Moth Pheromones. Causes and Consequences. Pp. 25–39 in R. T. Carde and J. D. Allison, eds. Pheromone Communication in Moths: Evolution, Behavior, and Application. University of California Press, Berkeley.

Conner, W. E., T. Eisner, R. K. V. Meer, A. Guerrero, and J. Meinwald. 1981. Precopulatory sexual interaction in an arctiid moth Utetheisa ornatrix: role of a pheromone derived from dietary alkaloids. Behav. Ecol. Sociobiol. 9:227–235.

Coyne, J. A., and H. A. Orr. 2004. Speciation. Sinauer, Sunderland, MA.

Croat, T. B. 1969. Seasonal Flowering Behavior in Central Panama. Ann. Mo. Bot. Gard. 56:295–307. Missouri Botanical Garden Press.

Darragh, K., K. J. R. P. Byers, R. M. Merrill, W. O. McMillan, S. Schulz, and C. D. Jiggins. 2019. Male pheromone composition depends on larval but not adult diet in Heliconius melpomene. Ecol. Entomol., doi: 10.1111/een.12716.

Darragh, K., G. MontejoLKovacevich, K. M. Kozak, C. R. Morrison, C. M. E. Figueiredo, J. S. Ready, C. Salazar, M. Linares, K. J. R. P. Byers, R. M. Merrill, W. O. McMillan, S. Schulz, and C. D. Jiggins. 2020. Species specificity and intraspecific variation in the chemical profiles of Heliconius butterflies across a large geographic range. Ecol. Evol. 10:3895–3918.

Darragh, K., A. Orteu, D. Black, K. J. R. P. Byers, D. Szczerbowski, I. A. Warren, P. Rastas, A. Pinharanda, J. W. Davey, S. F. Garza, D. A. Almeida, R. M. Merrill, W. O. McMillan, S. Schulz, and C. D. Jiggins. 2021. A novel terpene synthase controls differences in anti-aphrodisiac pheromone production between closely related Heliconius butterflies. PLOS Biol. 19:e3001022.

Darragh, K., S. Vanjari, F. Mann, M. F. Gonzalez-Rojas, C. R. Morrison, C. Salazar, C. Pardo-Diaz, R. M. Merrill, W. O. McMillan, S. Schulz, and C. D. Jiggins. 2017. Male sex pheromone components in Heliconius butterflies released by the androconia affect female choice. PeerJ 5:e3953.

De Cáceres, M., L. Coll, P. Legendre, R. B. Allen, S. K. Wiser, M.-J. Fortin, R. Condit, and S. Hubbell. 2019. Trajectory analysis in community ecology. Ecol. Monogr. 89:e01350.

De Cáceres, M., P. Legendre, S. K. Wiser, and L. Brotons. 2012. Using species combinations in indicator value analyses. Methods Ecol. Evol. 3:973–982.

Dray, S., and A. B. Dufour. 2007. The ade4 package: implementing the duality diagram for ecologists. J. Stat. Softw. 22:1–20.

Dressler, R. L. 1982. Biology of the Orchid Bees (Euglossini). Annu. Rev. Ecol. Syst. 13:373–394.

Eltz, T., C. Bause, K. Hund, J. J. G. Quezada-Euan, and T. Pokorny. 2015. Correlates of perfume load in male orchid bees. Chemoecology 25:193–199.

Eltz, T., D. W. Roubik, and K. Lunau. 2005a. Experience-dependent choices ensure species-specific fragrance accumulation in male orchid bees. Behav. Ecol. Sociobiol. 59:149.

Eltz, T., A. Sager, and K. Lunau. 2005b. Juggling with volatiles: exposure of perfumes by displaying male orchid bees. J. Comp. Physiol. A 191:575–581.

Eltz, T., W. M. Whitten, D. W. Roubik, and K. E. Linsenmair. 1999. Fragrance Collection, Storage, and Accumulation by Individual Male Orchid Bees. J. Chem. Ecol. 25:157– 176.

Eltz, T., Y. Zimmermann, J. Haftmann, R. Twele, W. Francke, J. J. G. Quezada-Euan, and K. Lunau. 2007. Enfleurage, lipid recycling and the origin of perfume collection in orchid bees. Proc. R. Soc. B Biol. Sci. 274:2843–2848. Royal Society.

Fournier, L. A., and S. Salas. 1966. Algunas observaciones sobre la dinámica de la floración en el bosque tropical húmedo de Villa Colón. Rev. Biol. Trop. 14:75–85.

Frankie, G. W., H. G. Baker, and P. A. Opler. 1974. Comparative Phenological Studies of Trees in Tropical Wet and Dry Forests in the Lowlands of Costa Rica. J. Ecol. 62:881–919. [Wiley, British Ecological Society].

Gerhardt, H. C. 1982. Sound Pattern Recognition in Some North American Treefrogs (Anura: Hylidae): Implications for Mate Choice. Integr. Comp. Biol. 22:581–595.

Groot, A. T., T. Dekker, and D. G. Heckel. 2016. The Genetic Basis of Pheromone Evolution in Moths. Annu. Rev. Entomol. 61:null.

Hervé, M. 2021. RVAideMemoire: Testing and Plotting Procedures for Biostatistics. R package version 0.9-81.

Janzen, D. H. 1967. Synchronization of Sexual Reproduction of Trees Within the Dry Season in Central America. Evolution 21:620–637. [Society for the Study of Evolution, Wiley].

Jiggins, C. D., R. E. Naisbit, R. L. Coe, and J. Mallet. 2001. Reproductive isolation caused by colour pattern mimicry. Nature 411:302–305.

Johansson, B. G., and T. M. Jones. 2007. The role of chemical communication in mate choice. Biol. Rev. 82:265–289.

Kassambara, A. 2019. ggpubr: “ggplot2” Based Publication Ready Plots. R package version 0.2.4. https://CRAN.R-project.org/package=ggpubr.

Liénard, M. A., M. Strandh, E. Hedenström, T. Johansson, and C. Löfstedt. 2008. Key biosynthetic gene subfamily recruited for pheromone production prior to the extensive radiation of Lepidoptera. BMC Evol. Biol. 8:270.

Löfstedt, C. 1993. Moth Pheromone Genetics and Evolution. Philos. Trans. R. Soc. Lond. B Biol. Sci. 340:167–177.

Martin, M. D., and T. C. Mendelson. 2016. The accumulation of reproductive isolation in early stages of divergence supports a role for sexual selection. J. Evol. Biol. 29:676– 689.

Mas, F., and J.-M. Jallon. 2005. Sexual Isolation and Cuticular Hydrocarbon Differences between Drosophila santomea and Drosophila yakuba. J. Chem. Ecol. 31:2747–2752.

McLean, M., D. Mouillot, M. Lindegren, S. Villéger, G. Engelhard, J. Murgier, and A. Auber. 2019. Fish communities diverge in species but converge in traits over three decades of warming. Glob. Change Biol. 25:3972–3984.

McPeek, M. A., L. B. Symes, D. M. Zong, and C. L. McPeek. 2011. Species recognition and patterns of population variation in the reproductive structures of a damselfly genus. Evol. Int. J. Org. Evol. 65:419–428.

Müller, K., and H. Wickham. 2022. tibble: Simple Data Frames.

Oksanen, J., F. Guillaume Blanchet, M. Friendly, R. Kindt, P. Legendre, D. McGlinn, P. Minchin, R. O’Hara, G. Simpson, P. Solymos, H. Stevens, E. Szoecs, and H. Wagner. 2020. vegan: Community Ecology Package. R package version 2.5-7.

Pemberton, R. W., and G. S. Wheeler. 2006. Orchid bees don’t need orchids: evidence from the naturalization of an orchid bee in Florida. Ecology 87:1995–2001.

Pfennig, K. S. 1998. The evolution of mate choice and the potential for conflict between species and mate–quality recognition. Proc. R. Soc. Lond. B Biol. Sci. 265:1743– 1748.

Pokorny, T., M. Hannibal, J. J. G. Quezada-Euan, E. Hedenström, N. Sjöberg, J. Bång, and T. Eltz. 2013. Acquisition of species-specific perfume blends: influence of habitat-dependent compound availability on odour choices of male orchid bees (Euglossa spp.). Oecologia 172:417–425.

R Core Team. 2021. R: A language and environment for statistical computing. R Foundation for Statistical Computing, Vienna, Austria.

Ramírez, S. R. 2019. Pollinator specificity and seasonal patterns in the euglossine bee-orchid mutualism at La Gamba Biological Station. Acta ZooBot Austria 156:171–181.

Ramírez, S. R., R. L. Dressler, and M. Ospina. 2002. Abejas euglosinas (Hymenoptera: Apidae) de la Región Neotropical: Listado de especies con notas sobre su biología. Biota Colomb. 3:7–118.

Ramírez, S. R., T. Eltz, F. Fritzsch, R. Pemberton, E. G. Pringle, and N. D. Tsutsui. 2010a. Intraspecific Geographic Variation of Fragrances Acquired by Orchid Bees in Native and Introduced Populations. J. Chem. Ecol. 36:873–884.

Ramírez, S. R., T. Eltz, M. K. Fujiwara, G. Gerlach, B. Goldman-Huertas, N. D. Tsutsui, and N. E. Pierce. 2011. Asynchronous Diversification in a Specialized Plant-Pollinator Mutualism. Science 333:1742–1746.

Ramírez, S. R., D. W. Roubik, C. Skov, and N. E. Pierce. 2010b. Phylogeny, diversification patterns and historical biogeography of euglossine orchid bees (Hymenoptera: Apidae). Biol. J. Linn. Soc. 100:552–572.

Roberts, D. R., W. D. Alecrim, J. M. Heller, S. R. Ehrhardt, and J. B. Lima. 1982. Male Eufriesia purpurata, a DDT-collecting euglossine bee in Brazil. Nature 297:62–63. Nature Publishing Group.

Robertson, H. M. 2019. Molecular Evolution of the Major Arthropod Chemoreceptor Gene Families. Annu. Rev. Entomol. 64:227–242.

Roelofs, W. L., and A. P. Rooney. 2003. Molecular genetics and evolution of pheromone biosynthesis in Lepidoptera. Proc. Natl. Acad. Sci. U. S. A. 100:9179–9184.

Ryan, M. J., and M. A. Guerra. 2014. The mechanism of sound production in túngara frogs and its role in sexual selection and speciation. Curr. Opin. Neurobiol. 28:54–59.

Saveer, A. M., P. G. Becher, G. Birgersson, B. S. Hansson, P. Witzgall, and M. Bengtsson. 2014. Mate recognition and reproductive isolation in the sibling species Spodoptera littoralis and Spodoptera litura. Chem. Ecol. 2:18.

Schneider, D. 1992. 100 years of pheromone research. An essay on lepidoptera. Naturwissenschaften 79:241–250.

Shahandeh, M. P., A. Pischedda, and T. L. Turner. 2018. Male mate choice via cuticular hydrocarbon pheromones drives reproductive isolation between Drosophila species. Evolution 72:123–135.

Smadja, C. M., and R. K. Butlin. 2008. On the scent of speciation: the chemosensory system and its role in premating isolation. Heredity 102:77–97.

Sturbois, A., M. D. Cáceres, M. Sánchez-Pinillos, G. Schaal, O. Gauthier, P. L. Mao, A. Ponsero, and N. Desroy. 2021. Extending community trajectory analysis: New metrics and representation. Ecol. Model. 440:109400.

Thioulouse, J., S. Dray, A.-B. Dufour, A. Siberchicot, T. Jombart, and S. Pavoine. 2018. Multivariate Analysis of Ecological Data with ade4. Springer.

Venables, W. N., B. D. Ripley, and W. N. Venables. 2002. Modern applied statistics with S.

Vogel, S. 1966. Parfümsammelnde Bienen als Bestäuber von Orchidaceen und Gloxinia. Österr. Bot. Z. 113:302–361. Springer.

Wang, Y., U. Naumann, S. T. Wright, and D. I. Warton. 2012. mvabund– an R package for model-based analysis of multivariate abundance data. Methods Ecol. Evol. 3:471– 474.

Warton, D. I., S. T. Wright, and Y. Wang. 2012. Distance-based multivariate analyses confound location and dispersion effects. Methods Ecol. Evol. 3:89–101.

Weber, M. G., L. Mitko, T. Eltz, and S. R. Ramírez. 2016. Macroevolution of perfume signalling in orchid bees. Ecol. Lett. 19:1314–1323.

Wei, T., and V. Simko. 2021. corrplot: Visualization of a correlation matrix.

West, R. J. D., and A. Kodric-Brown. 2015. Mate Choice by Both Sexes Maintains Reproductive Isolation in a Species Flock of Pupfish (Cyprinodon spp) in the Bahamas. Ethology 121:793–800.

Whitten, W., A. Young, and D. Stern. 1993. Nonfloral sources of chemicals that attract male euglossine bees (Apidae: Euglossini). J. Chem. Ecol. 19:3017–27.

Wickham, H. 2009. ggplot2: Elegant Graphics for Data Analysis. Springer-Verlag New York.

Wickham, H., R. François, L. Henry, and K. Müller. 2021. dplyr: A Grammar of Data Manipulation. R package version 1.0.7.

Wilke, C. O. 2020. cowplot: Streamlined Plot Theme and Plot Annotations for “ggplot2.”

Wyatt, T. D. 2014. Pheromones and Animal Behavior: Chemical Signals and Signatures. Cambridge University Press, Cambridge.

Wyatt, T. D. 2003. Pheromones and Animal Behaviour: Communication by Smell and Taste. Cambridge University Press, Cambridge.

Zimmermann, Y., S. R. Ramírez, and T. Eltz. 2009. Chemical niche differentiation among sympatric species of orchid bees. Ecology 90:2994–3008.

Zimmermann, Y., D. W. Roubik, and T. Eltz. 2006. Species-specific attraction to pheromonal analogues in orchid bees. Behav. Ecol. Sociobiol. 60:833–843.

