## Supplementary information for "Seasonal stability and species specificity of environmentally acquired chemical mating signals in orchid bees"

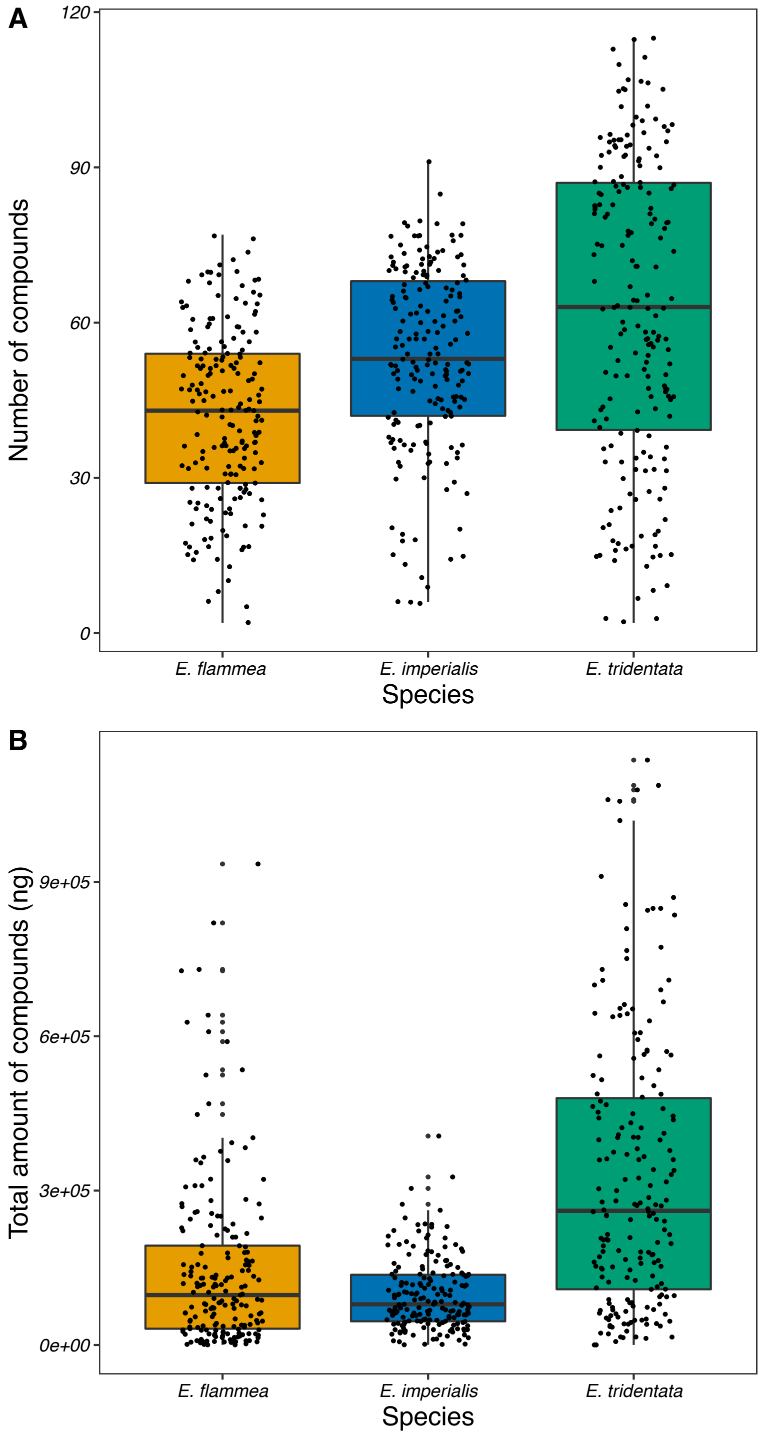


**Figure S1**: A) Species differ significantly in number of chemical compounds (ANOVA, $F_{2,569} = 40.06$, $p < .0001$). All pairwise test significantly different (post-hoc Tukey test p<0.001). B) Species differ significantly in the total amount of chemical compounds (ANOVA, $F_{2,569} = 80.48$, $p < .0001$). All pairwise test significantly different (post-hoc Tukey test p<0.05).

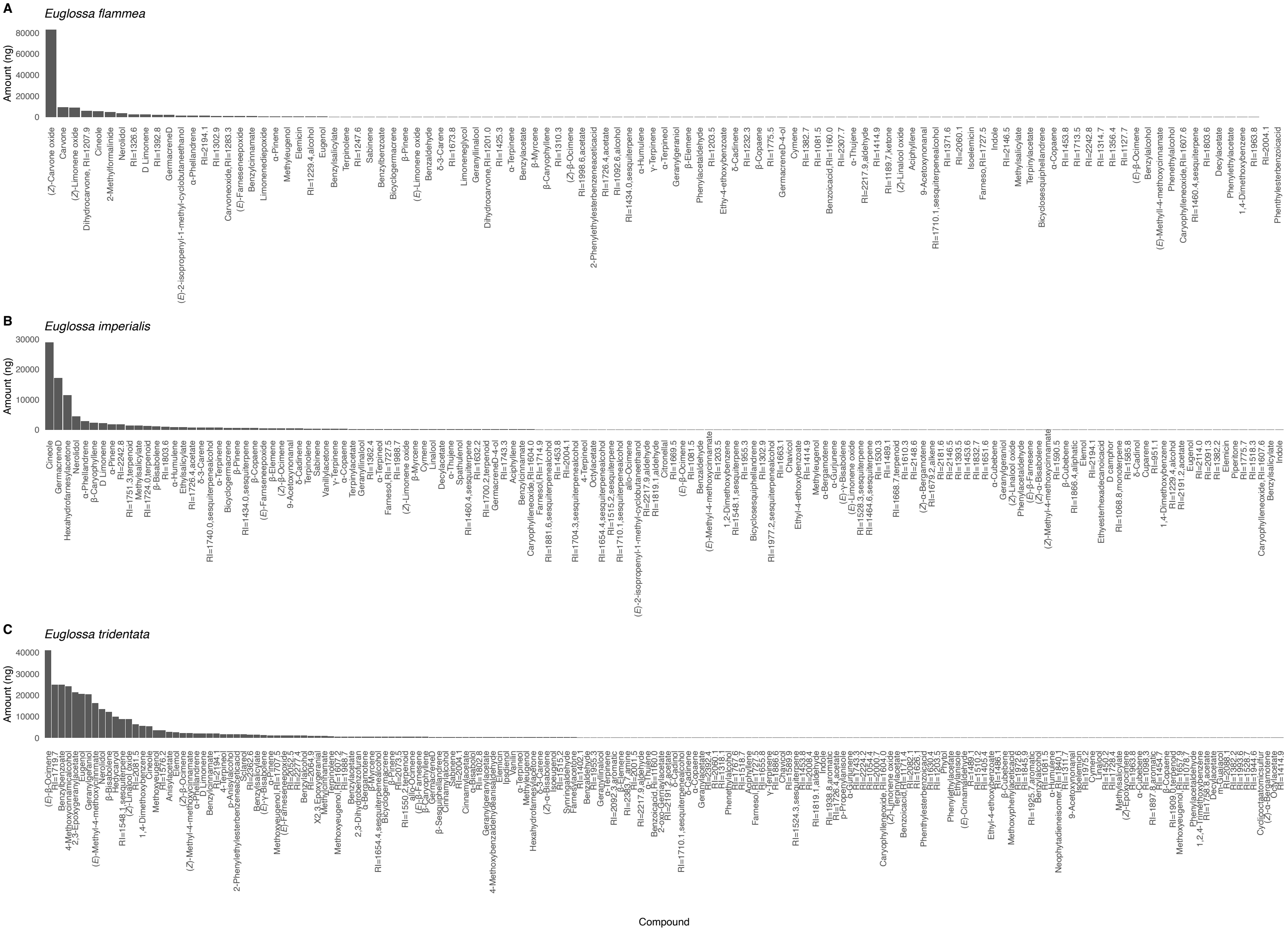


**Figure S2:** Barplots showing the average abundance (ng) of compounds in each species

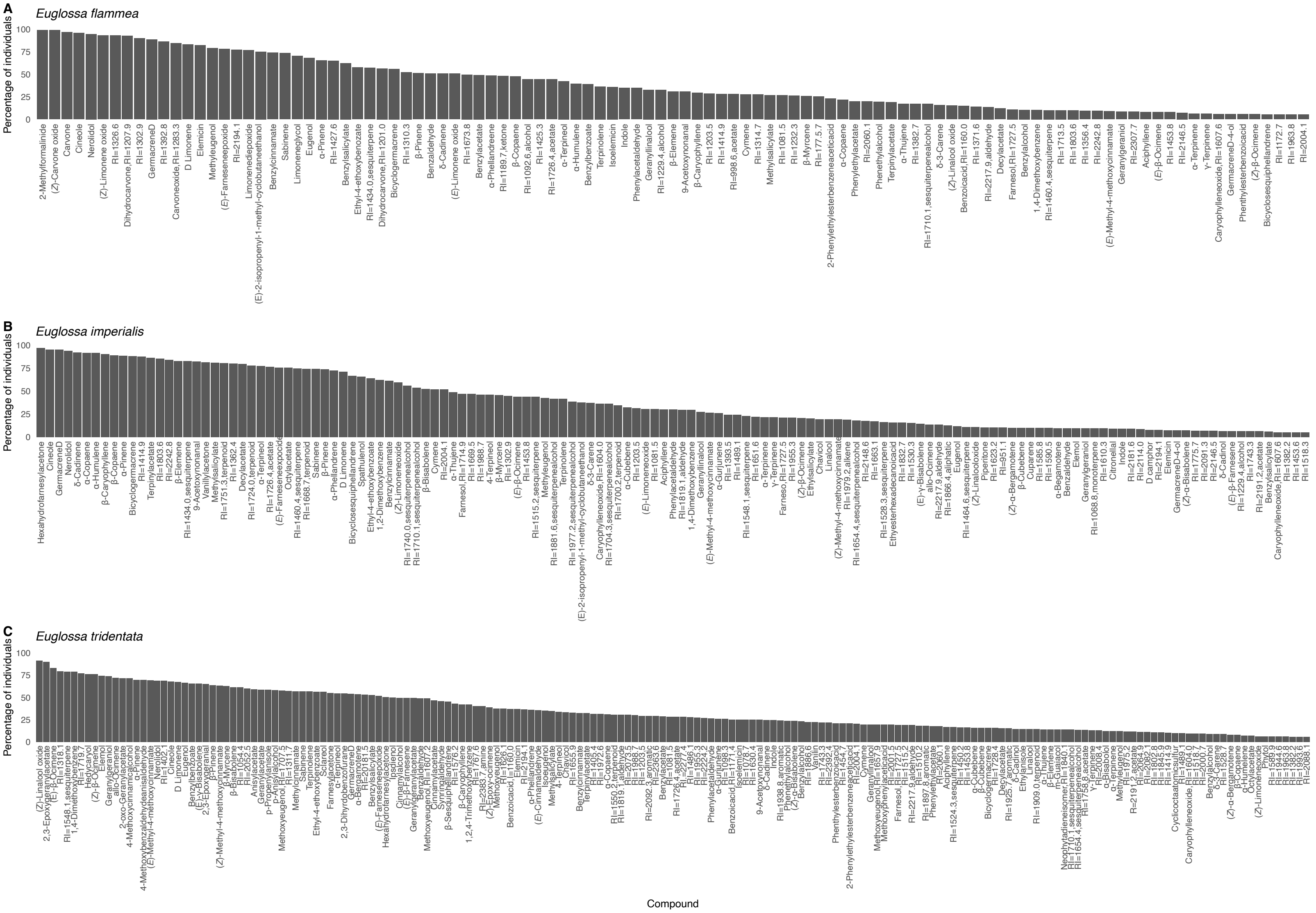

**Figure S3:** Barplots showing the frequency of compounds in each species.

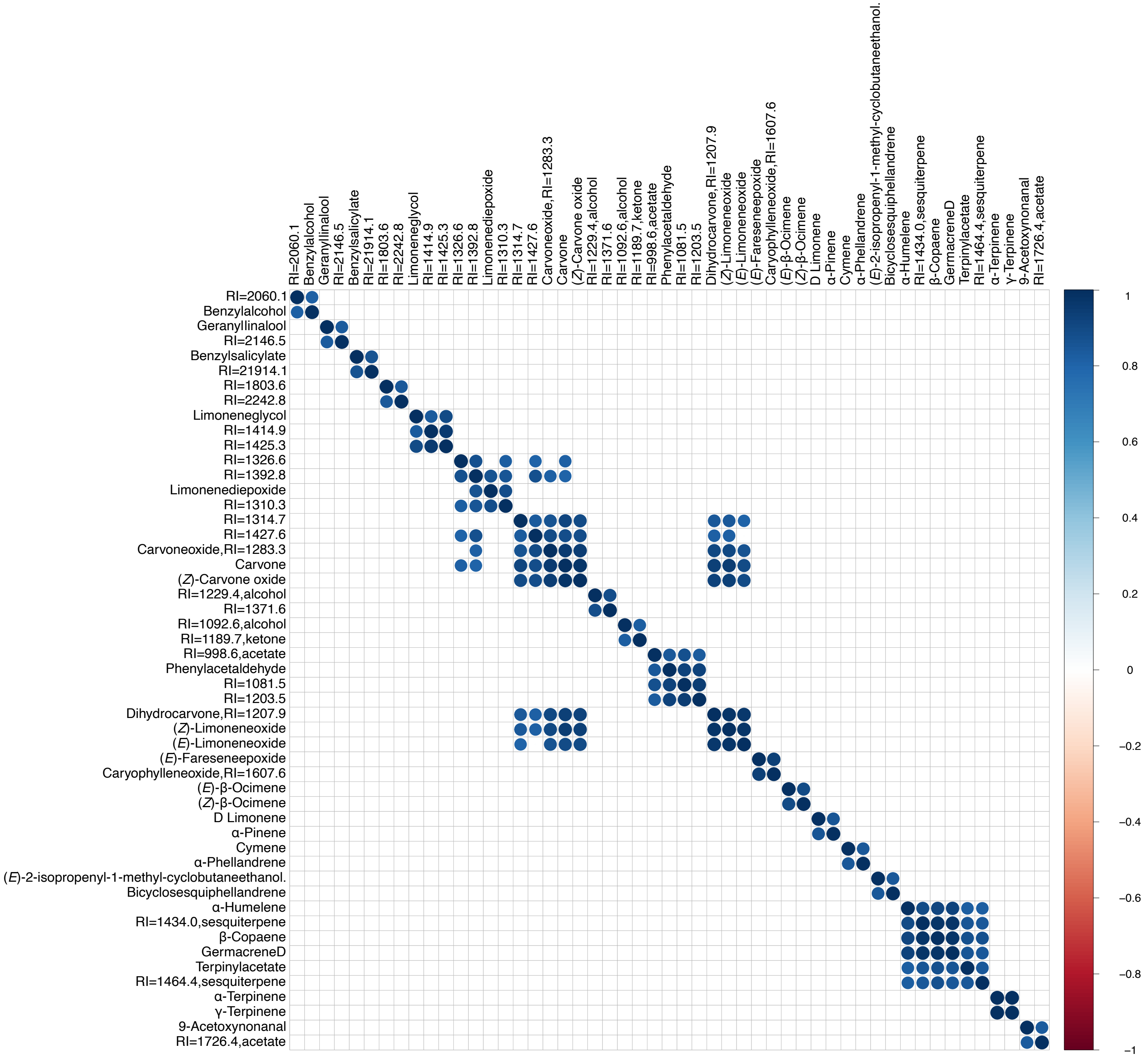


**Figure S4**: Sets of highly intercorrelated compounds (“motifs”) found in *E. flammea*. Compounds were considered highly correlated and therefore included if R>0.8 and p<0.01.

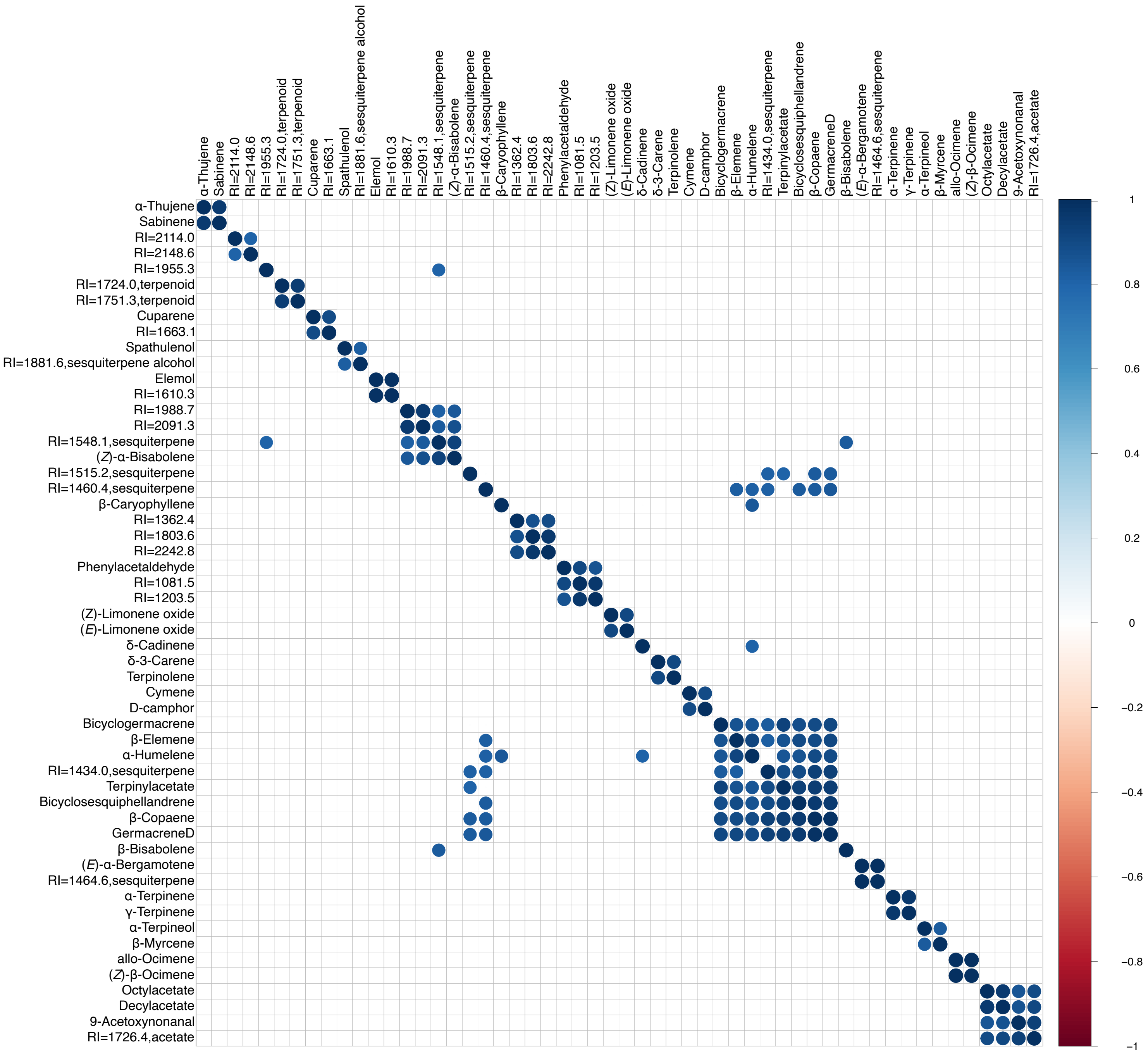


**Figure S5**: Sets of highly intercorrelated compounds (“motifs”) found in *E.imperialis*. Compounds were considered highly correlated and therefore included if R>0.8 and p<0.01.

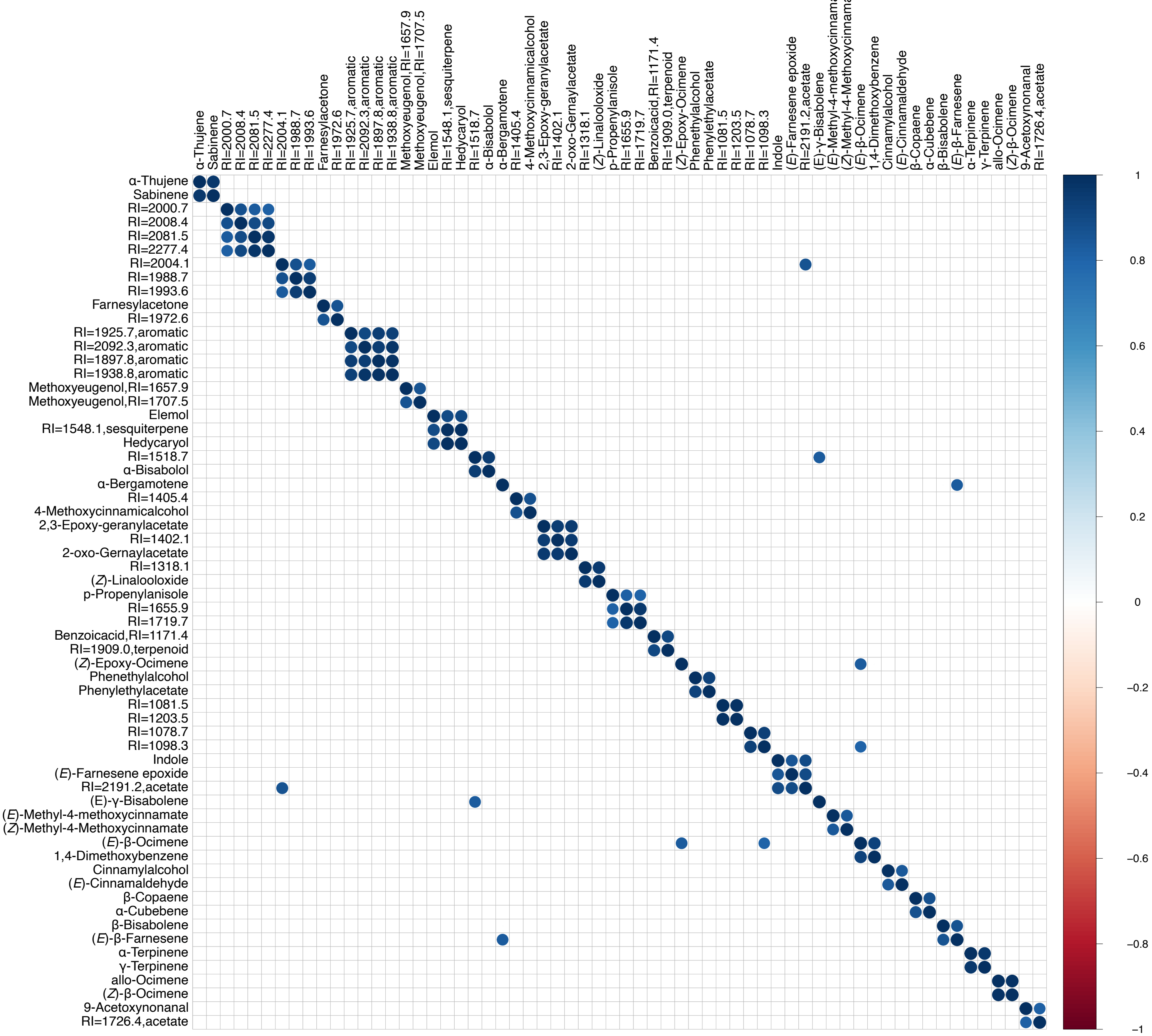


**Figure S6**: Sets of highly intercorrelated compounds (“motifs”) found in *E. tridentata*. Compounds were considered highly correlated and therefore included if R>0.8 and p<0.01.

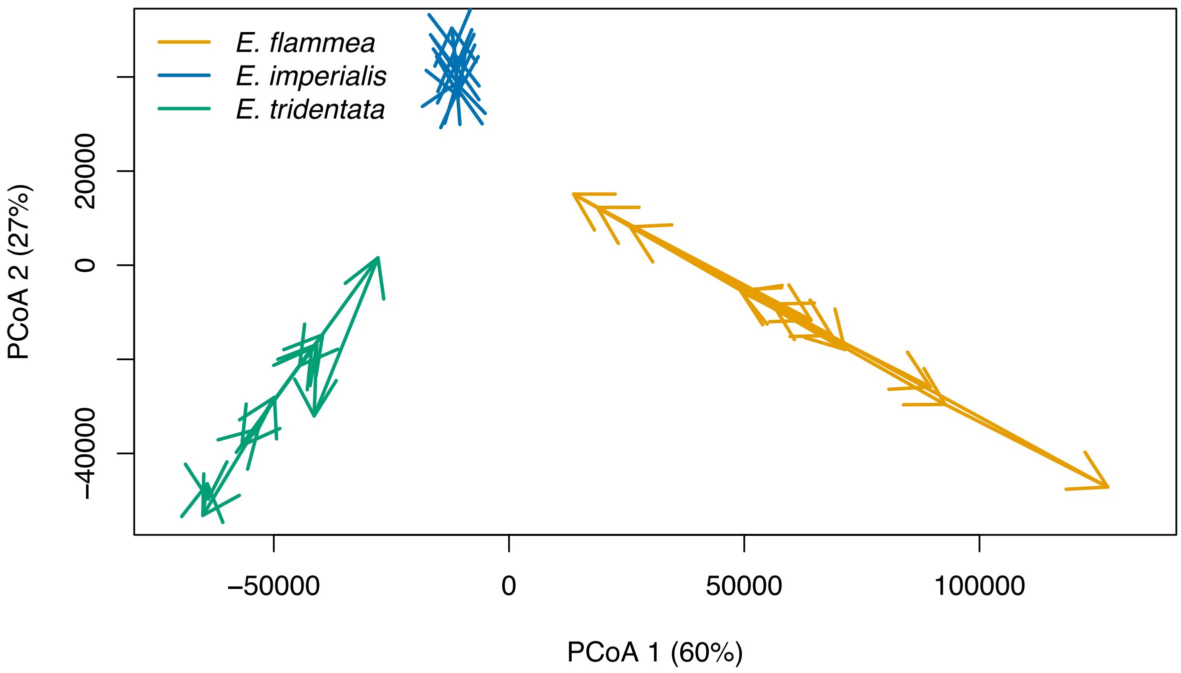


**Figure S7**: Community trajectory analysis of three orchid bee species throughout the study period of one year. Overall trajectory length is lower in *E. imperialis* (170,281), than for *E. flammea* and *E. tridentata* (551,225 and 590,910, respectively). All three species had low levels of overall directionality (*E. flammea*, 0.305; *E. imperialis*, 0.326; *E. tridentata*, 0.342).

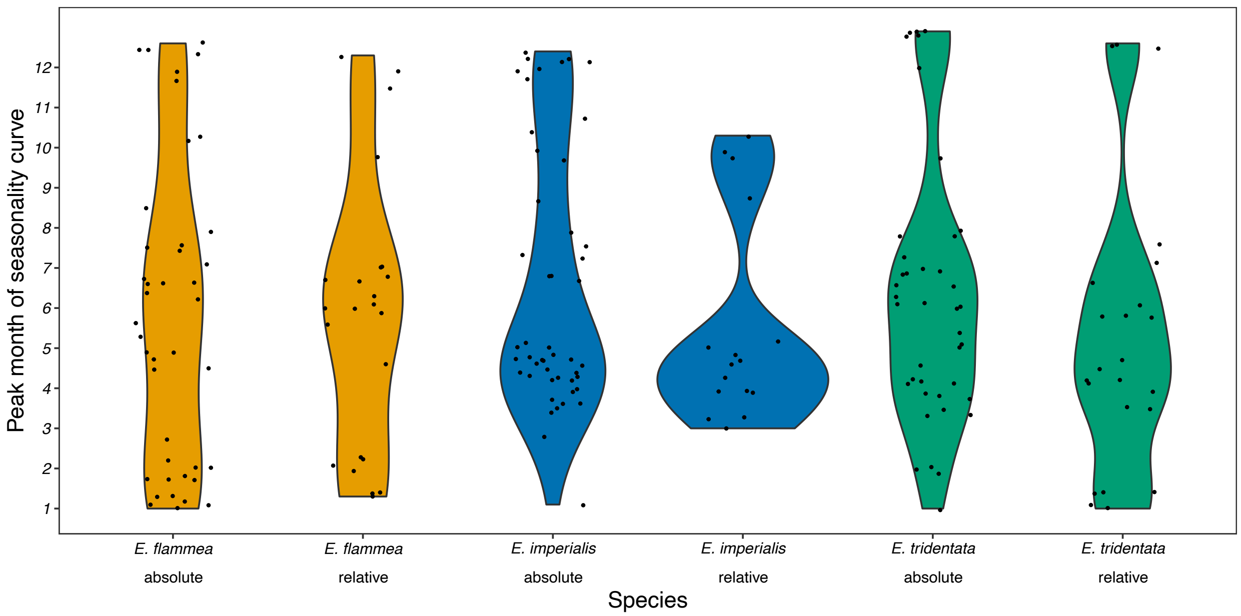


**Figure S8**: Violin plots illustrating the variation in the peak month of the seasonality curve for compounds of each species including both absolute and relative amounts. Only compounds which were determined to exhibit seasonality were included. (Kruskal Wallis, df=5, *H* test statistic=3.49, p=NS). For the y-axis, 1 is January and 12 is December.

**Table S1.** Model selection table for PERMANOVA models based on AIC scores

| Model | Residual Sum of Squares | DF | AIC |
|---|---|---|---|
| Chemical profile ~ Species + Month +Species*Month | 120.88 | 36 | 2820.62 |
| **Chemical profile ~ Species + Month** | **131.41** | **14** | **2820.40** |
| Chemical profile ~ Species | 137.90 | 2 | 2823.96 |

Note: Best fit models are highlighted in bold.

**Table S2.** Analysis of deviance model selection table for multivariate generalized linear models based on likelihood ratio tests

| Model | ΔDeviance | Residual DF | *p*-value |
|---|---|---|---|
| Chemical profile ~ Species + Month +Species*Month | | 553 | |
| **Chemical profile ~ Species + Month** | **10073** | **557** | **0.089** |
| Chemical profile ~ Species | 11121 | 569 | 0.001 |

Note: Each model is compared to the model above it. Best fit models are highlighted in bold.

**Table S3.** Pairwise comparisons of dispersion of chemical profiles of *E. flammea, E. imperialis* and *E. tridentata* (Observed p-value below diagonal, permuted p-value above diagonal).

| | *E. flammea* | *E. imperialis* | *E. tridentata* |
|---|---|---|---|
| *E. flammea* | | 0.41 | 0.001 |
| *E. imperialis* | 0.46 | | 0.001 |
| *E. tridentata* | <0.0001 | <0.0001 | |

**Table S4.** Summary of ManyGLM results including all significant explanatory variables with data including all species across the entire sampling period.

| Parameter | Residual DF | DF | Deviance | *p*-value | Compounds |
|---|---|---|---|---|---|
| Species | 569 | 2 | 29786 | 0.001 | Unknown (RI=1318.1), vanillyl acetone, hexahydrofarnesylacetone*, *E*-p-methyl cinnamyl acetate, 4-methoxy cinnamic alcohol, 2-oxo-geranylacetate, 2,3-epoxygeranyl acetate*, unknown (RI=1402.1), hedycaryol, unknown (RI=1803.6)* |
| Month | 557 | 12 | 11121 | 0.001 | Ethyl 4-ethoxybenzoate, unknown (RI=1203.5), unknown acetate (RI=1081.5), 2-methylformalinide, dihydrocarvone (2), carvone oxide, phenyl acetaldehyde, geranylacetate, limonene diepoxide, unknown (RI=1427.6) |

Note: The compound list includes the ten compounds that contribute the most to the deviance explained by each variable. The compounds are listed in descending order of contribution to deviance. Compounds highlighted with * were also identified by the indicator analysis.

**Table S5.** Results of cosinor model analysis of average species' differences throughout the year to test for seasonality.

| Species | Phase | Cosine *p*-value | Sine *p*-value | Seasonal? |
|---|---|---|---|---|
| *E. flammea – E. imperialis* | 4.1 | 1 | 0.86 | FALSE |
| *E. flammea – E. tridentata* | 6.6 | 0.19 | 1 | FALSE |
| *E. imperialis – E. tridentata* | 5.9 | 0.19 | 0.62 | FALSE |

**Note: alpha threshold set at 0.025**

**Table S6.** Results of cosinor model analysis of species' NMDS dimensions throughout the year to test for seasonality of overall perfume variation.

| Species | NMDS dimension | Phase | Cosine *p*-value | Sine *p*-value | Seasonal? |
|---|---|---|---|---|---|
| *E. flammea* | MDS1 | 12.6 | <0.0001 | 1 | TRUE |
| *E. flammea* | MDS2 | 1.1 | 0.055 | 1 | FALSE |
| *E. imperialis* | MDS1 | 4.7 | 1 | 0.037 | FALSE |
| *E. imperialis* | MDS2 | 7.8 | 0.2 | 1 | FALSE |
| *E. tridentata* | MDS1 | 12 | <0.0001 | 0.012 | TRUE |
| *E. tridentata* | MDS2 | 6.5 | <0.0001 | 0.26 | TRUE |
| *E. tridentata* | MDS3 | 3 | 1 | 0.17 | FALSE |

**Note: alpha threshold set at 0.025**

**Table S7.** Results of cosinor model analysis of number and amount of compounds collected by each species throughout the year to test for seasonality.

| Species | Characteristic | Phase | Cosine *p*-value | Sine *p*-value | Seasonal? |
|---|---|---|---|---|---|
| *E. flammea* | No. of compounds | 3.4 | 0.61 | 0.13 | FALSE |
| *E. imperialis* | No. of compounds | 3.4 | 0.60 | 0.055 | FALSE |
| *E. tridentata* | No. of compounds | 5 | 0.29 | 0.058 | FALSE |
| *E. flammea* | Amount (ng) | 1.4 | 0.027 | 0.61 | FALSE |
| *E. imperialis* | Amount (ng) | 7.1 | 0.30 | 0.96 | FALSE |
| *E. tridentata* | Amount (ng) | 6.3 | 0.0012 | 0.19 | TRUE |

**Note: alpha threshold set at 0.025**
